# Generating Hybrid Proteins with the MSA-Transformer

**DOI:** 10.1101/2025.11.20.689447

**Authors:** Sanjana Tule, Samuel Davis, Ivan Koludarov, Ariane Mora, Mikael Bodén

## Abstract

Protein superfamilies display extensive sequence and functional divergence, providing a rich landscape for engineering functionally enhanced variants. We present a stochastic, iterative framework that leverages the MSA-Transformer to generate intermediate sequences between a homologous pair of user-specified “source” and “target” proteins in sequence space. Sequence sites to “mask” is either selected by embedding-based dissimilarity or by row-attention information, while beam search concurrently explores multiple mutational pathways. Pre-trained sparse autoencoders combined with sequence and structural analyses are used to trace the inheritance and exchange of features across the “mutational pathway”, revealing “hybrid” sequences that integrate properties of both source and target proteins. Applied across diverse protein families, the framework produces sequences that occupy biologically meaningful regions of sequence space and achieve higher consistency and plausibility scores than random baselines according to *in silico* metrics. In the B1/B2 metallo-*β*-lactamase family, hybrids largely retain their core fold recombining structural and active-site motifs from both subclasses, demonstrating the model’s capacity to preserve catalytic features while exploring novel structural permutations.

**Code availability:** The implementation is available via GitHub at https://github.com/santule/protmixy.

## 1. Introduction

Proteins are the fundamental molecular engines of life, adapted over billions of years of evolution to diverse evolutionary pressures to perform an extraordinary range of structural and functional roles. Protein super-families arise from shared origins but typically display a broad spectrum of functional and sequence variation, while retaining largely conserved yet subtly variable structures [7, 20]. While closely related proteins tend to share similar sequences and functions, more distantly related members can encode distinct biochemical capabilities [14]. This functional heterogeneity distributed across a superfamily represents a rich reservoir for protein design [3]. Rather than engineering new proteins from a single template, the diversity within families provides an opportunity to *blend* properties from *multiple* proteins to generate entirely novel sequences with hybrid characteristics.

Ancestral sequences, reconstructed using phylogenetic trees and evolutionary models, represent inferred progenitors of modern proteins. They retain sequence-structure elements that have persisted despite subsequent divergence. Importantly, ancestral proteins already embody a natural form of hybridisation, when viewed from the perspective of present-day sequences: they reconcile traits from distinct evolutionary branches and integrate information from multiple lineages. This intrinsic composite nature often confers properties such as enhanced stability, functional breadth, or evolvability–making ancestral reconstructions both powerful tools for protein engineering and conceptual precursors to artificial hybrids [12, 29, 28, 37, 5, 36]. In this sense, ancestral proteins illustrate why generating hybrids is meaningful: evolution itself has repeatedly arrived at hybrid-like intermediates as efficient, versatile solutions within sequence space.

While ancestral sequence reconstruction relies on explicit phylogenetic models and curated evolutionary histories, recent advances in deep learning-based protein language models [22, 33, 10, 15] implicitly capture evolutionary regularities from large-scale sequence datasets, offering a complementary data-driven approach to explore evolutionary sequence space [16, 18, 8, 38, 6]. Protein language models have also been successfully applied across a broad range of protein-related tasks, including residue–residue contact prediction, mutational effect inference, and the prediction of diverse protein properties and functions [31, 27, 24]. Collectively, protein language models offer a useful framework for exploring protein sequence space and evaluating the plausibility of generated or unseen sequences.

Earlier protein engineering studies have explored the generation of hybrid proteins by recombining fragments from homologous sequences while preserving structural compatibility [39]. More recent studies have investigated machine learning–guided navigation of protein sequence landscapes to identify improved variants [41]. In parallel, deep generative models have been applied to protein design, including approaches that design proteins *de novo* or optimise existing sequences using structure-based objectives [25, 11, 17]. Other studies have investigated exploration of latent representations learned by large protein language models to analyse evolutionary and structural relationships between sequences [32, 23]. However, these approaches typically focus on generating sequences from a single starting point or sampling broadly within sequence space. In contrast, our work explicitly investigates the generation of hybrid proteins by iteratively traversing the sequence representation space between a specified source and target sequence within a protein family.

Here, we use a protein language model to generate hybrid proteins—novel sequences that combine attributes from homologous sequences which share a common evolutionary origin but diverged in sequence, structure, and function. Leveraging the generative power of MSA-Transformer [35, 19], we hypothesise that the internal representations learnt from millions of multiple sequence alignments enable the model to generate plausible hybrids by recombining compatible features from related proteins.

To realise this, we *simulate* mutational pathways from a *source* to a *target* sequence, using the MSA-Transformer to iteratively introduce substitutions that steer the source sequence toward the target. The intermediate sequences generated along these pathways represent the hybrid—variants that integrate sequence, structural, and functional characteristics of both endpoints. Given that the MSA-Transformer draws on both local alignment context and evolutionary knowledge acquired through large-scale pre-training [33], we expect that the model will guide mutations in ways that remain compatible with constraints observed within natural protein families.

Hybrid sequences are produced through iterative masking and resampling from the model’s conditional probability distribution, with convergence defined by high similarity to the target sequence. We aim to determine whether the MSA-Transformer, when provided with an appropriately defined MSA conditioning context, can be steered to generate coherent hybrid intermediates between homologous proteins. In particular, we investigate how performance varies across different levels of sequence identity between the source and target sequence.

To evaluate the quality and plausibility of generated intermediates, we establish a set of *in silico* scoring metrics that assess whether the generated hybrids remain compatible with constraints observed in natural protein families. These include sequence and structural similarity to the source and target proteins, plausibility scores derived from established protein language models, and comparisons against random mutation baselines. Finally, we analyse the geometry of the mutational pathways in latent space, asking whether the hybrids lie along a simple linear interpolation between source and target sequence, or instead occupy non-linear regions of the representation space.

## 2. Results

### 2.1 Context and sampling steer MSA-Transformer mutational pathways

We first ask whether the composition of the input MSA can be exploited to steer the MSA Transformer to generate variants that form a mutational pathway between a source sequence (*S*) and a target sequence (*T*). Here, steering refers to conditioning the model on a curated MSA (*N*) constructed specifically for a given source–target pair. The curated MSA *N* along with *S* and *T* serves as input to the MSA-Transformer. We refer to *N* as the *conditioning context* under which it becomes conducive for the model to generate a mutational pathway connecting *S* and *T*. The mutational pathway generation begins by masking residues in *S* and decoding the most probable sequence (*C*) from the model’s output probability distribution (see Supplementary Methods, Section S1.1). The probabilistic acceptance criterion then determines whether to retain or discard the candidate sequence *C* based on the cosine distance to *T*. We use cosine distance in the embedding space as the optimisation objective, which we found empirically to yield more consistent convergence of mutational pathways than Euclidean distance (see Supplementary Methods). If retained, *C* replaces *S* as a new source sequence for the next iteration; otherwise, *S* is retained for subsequent iterations. The iterations stop when convergence criteria is met (see Figure 1A and Supplementary Methods, Section S1.2 for more details).

**Fig. 1.**
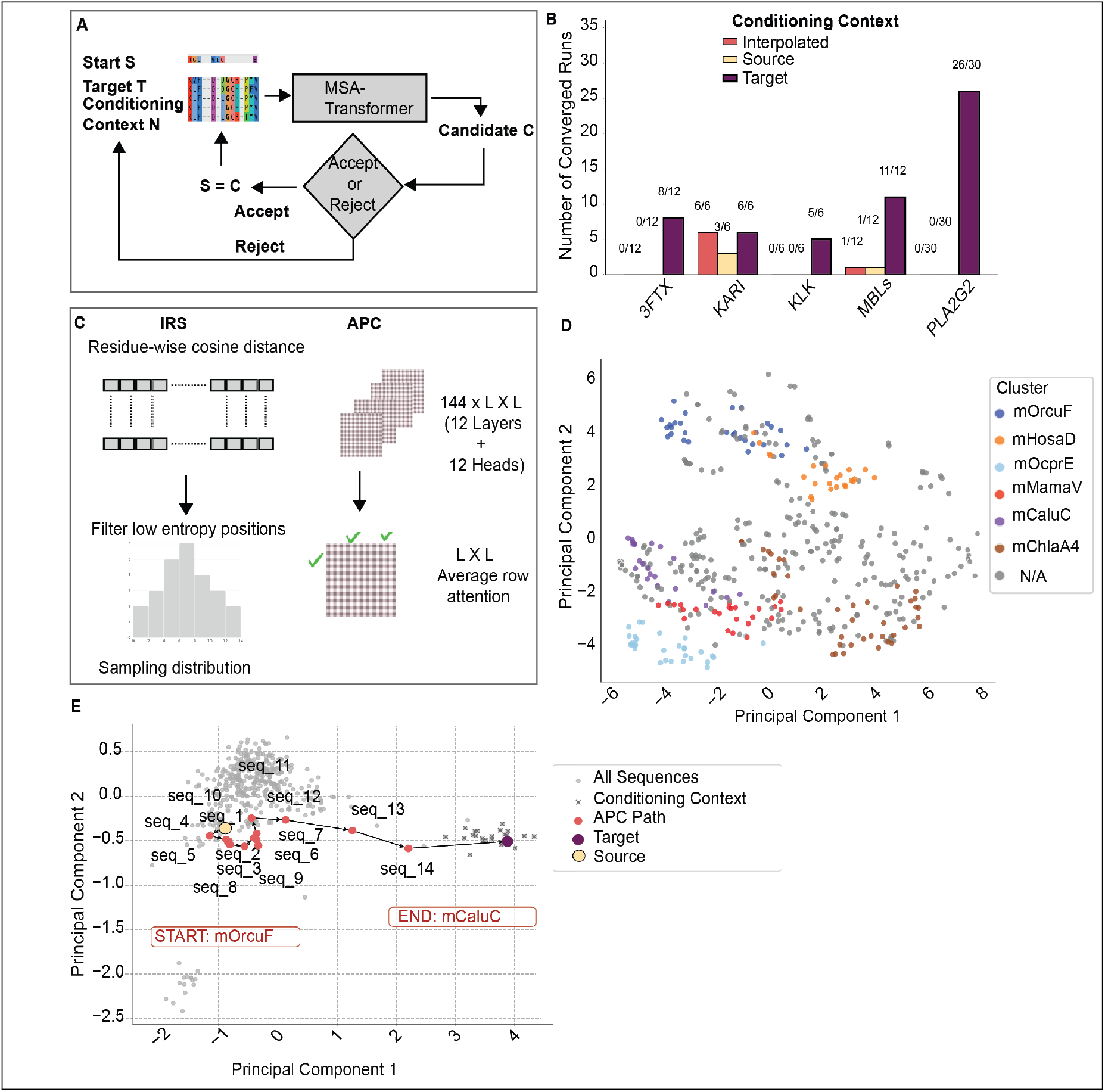
Target-conditioning context steers the MSA Transformer to generate mutational pathway between homologous source and target sequences. (A) Schematic overview of the mutational pathway generation framework, illustrating how hybrid sequences are iteratively generated from a source sequence (*S*) toward a target sequence (*T*) through intermediate candidate (*C*) sequences. (B) Convergence of sequences toward target sequences under different conditioning contexts (Target, Start, and Interpolated) for APC across multiple protein families. (C) Masking strategies used in the mutational pathway generation: IRS and APC. (D) Sequence clusters identified using the MSA-Transformer embeddings for the PLA2G2 family, visualised in the PCA Space. Cluster representatives are labelled, with clusters shown in distinct colours. Sequences not belonging to any clusters is shown as N/A. (E) Mutational pathway from *S* (*mOrcuF*) to *T* (*mCaluc*) within the PLA2G2 family, visualised in PCA space using APC masking strategy. PCA components were derived from ESM2 embeddings [23] computed for all sequences in the protein family’s MSA. Generated and input sequences were projected using the same PCA model. Input sequences are shown as grey circles, Target-focused context sequences as grey crosses, the source and target sequences in red, and intermediate sequences along the pathway in green. Arrows indicate the order of mutations.

The conditioning context *N* is constructed by clustering sequences within each protein family using a combination of HDBSCAN [26] and k-nearest neighbour (KNN). The sequence closest in cosine distance to the mean MSA Transformer–derived sequence embedding of each cluster is selected as the cluster representative, and these representatives serve as potential source (*S*) and target (*T*) sequences in different combinations. Figure 1D illustrates the clustered sequence space for a representative protein family, with distinct regions shown in different colours and cluster representatives labelled accordingly.

To evaluate the role of context, we test three conditioning contexts.

- Target-conditioning context: *N* comprised of sequences from the same cluster as the target sequence *T*.
- Start-conditioning context: *N* comprised of sequences from the same cluster as the start sequence *S*.
- Interpolated-conditioning context: *N* was populated with interpolated sequences between the start *S* and the target sequence *T*, in addition to the Target-conditioning context.

We evaluated the above settings across five diverse protein families: ketol-acid reductoisomerase (KARI, Class I), an essential enzyme in branched-chain amino acid biosynthesis; phospholipase A2 group 2 (PLA2G2), a secreted enzyme involved in lipid hydrolysis and inflammation as well as in viper snakes’ envenomation; B1/B2 metallo-*β*-lactamases (MBLs), antibiotic-resistance enzymes with broad substrate specificity; three-finger toxins (3FTx), a family of small, disulfide-rich proteins found in snake venoms; and tissue kallikreins (KLK), serine proteases with diverse physiological and pathological roles. For each family, **cluster representatives** are selected as *S* and *T* to define distinct sequence pairs. The number of clusters varied across families, leading to different total combinations of representative pairs: 12 pairs for 3FTX, 6 for KARI, 6 for KLK, 12 for MBLs, and 30 for PLA2G2.

We also examined how different residue masking strategies influence the model’s ability to generate mutational pathways. We compared two approaches. The first, independent residue sampling (IRS), prioritises residues that differ most from the target based on their cosine distance. The second, attention position coupling (APC), extends this strategy by incorporating row-attention information to account for relationships between positions learnt by the model. A schematic of the approach is shown in Figure 1C, with further details provided in the Supplementary Methods, Section S1.3.

All simulations were conducted using the same input MSAs, ensuring that the observed differences could be attributed to the MSA-based settings rather than stochastic variation in the mutational pathway generation process.

The results highlight that the conditioning context plays a critical role in generating the mutational pathways from *S* to *T*. In the Target-conditioning context, simulations consistently converged: starting sequences mutated toward the target sequences in the majority of runs (Figure 1B). In contrast, the Start- and Interpolated-conditioning contexts produced substantially lower or no convergence. The effect of the masking strategy (IRS vs. APC) on convergence was comparatively minor, with both approaches showing similar convergence trends across families (Figure 1B and Supplementary Figure S1). However, the pathways they followed differed. Figure 1E and Supplementary Figure S2 illustrates an example mutation pathways from *S* to *T* in PCA space, illustrating how IRS and APC explore sequence space differently.

### 2.2 Beam Search–Guided Non-Linear Mutational Pathways Improve Variant Quality Beyond Random Baselines

To better capture the diversity of possible routes between *S* and *T*, we extended our approach to explore multiple pathways in parallel. To achieve this, we incorporated a beam search strategy guided by the MSA-Transformer’s negative log-likelihood score and the cosine distance between the candidate sequence *C* and target sequence *T*. The former encourages sequences that remain plausible under the model, while the latter promotes directional progress towards *T* (see Supplementary Methods, Section S1.4).

To examine how sequence divergence shapes mutational pathways, we selected source (*S*) and target (*T*) sequences across a range of sequence identity thresholds. Using the same clusters defined in Results Section 2.1, for each cluster representative *T* in a protein family, a starting sequence *S* was randomly drawn from the remainder of the MSA such that its pairwise identity to *T* fell within a pre-defined range. Sequence identity was discretised into bins to systematically sample pairs with varying degrees of divergence. We repeated this procedure across all protein families and all clusters within each family (identified as per described in Supplementary Methods, Section S1.2), resulting in 14 sequence pairs for B1/B2 MBLs, 12 for KARI, 28 for PLA2G2, 11 for KLK, and 16 for 3FTx. The resulting distribution of sequence identity is shown in Supplementary Figure S3, comprising of 13 pairs in the 10-50% range, 10 in 50-60%, 20 in 60-70, 23 in 70-80% and 15 in 80-90%.

Under beam search, convergence rose from 54–62% in the 10–50% identity bin to 95–100% at 60–70%, then declined to 87–91% for 70–90% identity pairs. (Figure 2A). Although overall convergence counts were comparable between masking strategies, APC consistently achieved slightly higher convergence rates and required fewer iterations to reach convergence compared to IRS (Figure 2A; Supplementary Table S1 and S2). The improved performance of the attention-guided strategy is consistent with prior findings that row attention in the MSA-Transformer encodes inter-residue contact information [33], suggesting the attention based couplings capture informative residue dependencies that guide the search toward viable intermediates with fewer steps.

**Fig. 2.**
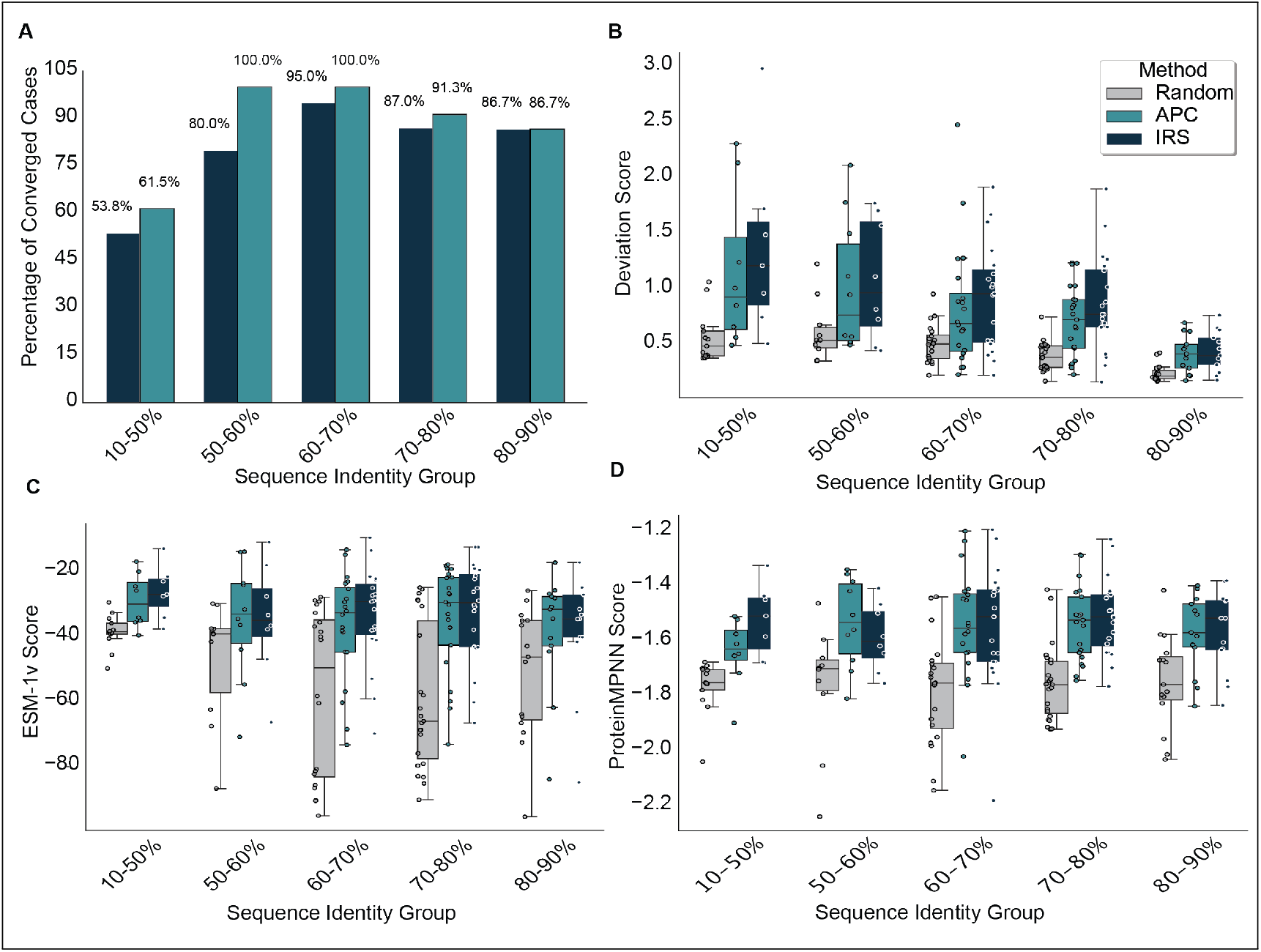
Beam search–guided non-linear mutational pathways improve variant quality beyond random baselines. (A) Convergence rates of beam search as a function of sequence identity between source (*S*) and target (*T*) sequences for the IRS (purple) and APC (yellow) generators. Both methods show improved convergence at higher sequence identities, with APC achieving slightly higher success rates overall and the highest convergence observed in the 60–80% range. (B) Deviation score from the linear interpolation path between the source (*S*) and target (*T*) embeddings, stratified by sequence identity. Lower deviation values indicate pathways that more closely follow the straight path in embedding space. Both IRS and APC produce significantly higher deviation scores than random baselines, particularly at intermediate sequence identities, suggesting smoother transitions through embedding space. (C) Distribution of ESM-1v variant scores for generated sequences, stratified by sequence identity and generator method. Both IRS and APC achieve higher variant scores relative to the random baseline, indicating preservation of functional plausibility across all identity groups. (D) Distribution of ProteinMPNN scores for generated sequences, stratified by sequence identity and generator method. Higher (less negative) scores indicate better compatibility between sequence and structure, with APC showing consistently favourable distributions compared to random sampling.

To characterise the geometry of the converged mutational pathways, we quantified the deviation of each pathway from the straight line connecting the source and target embeddings using the deviation score (see Supplementary Methods, Section S1.5). This metric measures how closely a generated pathway follows the linear interpolation between *S* and *T* in the ESM2 [23] embedding space (650M-parameter ESM2 model), thereby indicating whether the search progresses along a direct or curved route through representation space.

We performed this analysis using ESM2 embeddings rather than MSA-Transformer embeddings because ESM2 operates on single sequences, enabling a consistent comparison between model-generated pathways and randomly sampled sequences in the same representation space. The deviation score is used purely as a geometric diagnostic and is not part of the optimisation objective used during pathway generation. Consequently, deviations from the linear interpolation reflect the geometry of the embedding space explored by the model rather than being directly induced by the optimisation procedure. Importantly, higher deviation is not interpreted as intrinsically desirable; rather, the deviation score is used to determine whether generated pathways simply follow the direct interpolation between *S* and *T* or instead traverse alternative regions of the representation space.

As a baseline, we implemented a random generator that, at each iteration, replaced 5% of differing sites between *S* and *T* with the corresponding residues from *T*. For each sequence pair, the random process was repeated across 10 independent seeds. Deviation scores were then estimated by sampling 100 pathways per protein sequence pair per method and averaging the results to obtain a stable comparison baseline.

The random baseline is intentionally simple and serves a diagnostic purpose rather than a competitive one. In our experiments, it is used to contrast mutations suggested by the MSA-Transformer with unguided sequence changes. Stronger baselines–such as generation using protein language models without MSA conditioning–would provide a complementary evaluation and represents an interesting direction for future work.

Across all sequence identity groups, IRS and APC guided pathways exhibited significantly higher deviation scores than the random baseline (paired Wilcoxon signed-rank tests with Benjamini-Hochberg correction for multiple comparisons; *q <* 0.05 for all bins), indicating that model-guided pathways do not simply follow the direct interpolation between *S* and *T* (Figure 2B). The curved pathways generated via IRS and APC suggest that the iterative process follows structured pathways shaped by the organisation of the MSA-Transformer’s learnt representation space, rather than random or linear interpolation. In the 10–50% identity group, APC (mean = 1.16 vs. 0.57 for random; *q* = 0.0156, *r* = 0.94) and IRS (1.37 vs. 0.56; *q* = 0.026, *r* = 0.91) both showed substantial deviation from linearity, though this range contained fewer sequence pairs and lower overall convergence rates, limiting the statistical power and generality of these results. Similar trends were observed for 50–60% identity APC: *q* = 0.039, *r* = 0.70; IRS: *q* = 0.026, *r* = 0.85). Statistical significance was strongest for the 60–80% identity range, where both methods achieved large effect sizes (*r >* 0.9) and highly significant differences after multiple-testing correction (*q <* 0.01 in all cases). Even at high similarity (80–90%), APC (0.41 vs. 0.23, *q* = 0.00048, *r* = 1.00) and IRS (0.43 vs. 0.23, *q* = 0.00082, *r* = 0.97) remained significantly higher than chance (see Supplementary Table S3).

We next assessed the biological plausibility of generated sequences using ESM-1v and ProteinMPNN variant scores as recommended by [19], as these metrics were experimentally validated to correlate with functional enzyme variants. For each protein pair, 100 mutational pathways were sampled per method with replacement, and the mean of the top three variant scores was used to obtain a stable estimate of sequence quality. The same aggregated values were used for both visualisation (Figures 2C and 2D) and statistical testing (see Supplementary Tables S4 and S5).

Paired Wilcoxon signed-rank tests with Benjamini-Hochberg correction for multiple comparisons confirmed that both APC- and IRS-guided searches produced significantly higher ESM-1v and ProteinMPNN variant scores than random sampling across nearly all sequence-identity ranges (Figures 2C–D).

At low divergence (10–50 %), both methods yielded substantial improvements over random for ESM-1v (APC: *q* = 0.026, *r* = 0.85; IRS: *q* = 0.026, *r* = 0.91) and ProteinMPNN (APC: *q* = 0.0167, *r* = 0.94; IRS: *q* = 0.026, *r* = 0.91), although interpretation in this range is limited by smaller sample sizes (*n* = 6–8).

Improvements were also observed for the 50–60 % identity group (ESM-1v: APC *q* = 0.041, *r* = 0.70; IRS *q* = 0.073, ns; ProteinMPNN: APC *q* = 0.0293, *r* = 0.74), suggesting consistent albeit slightly reduced effects at this divergence level.

The most pronounced gains occurred at intermediate sequence identities (60–80%), where both metrics showed large effect sizes (*r >* 0.9) and highly significant differences after multiple-testing correction (*q <* 0.001 in all cases). In this range, mean ESM-1v scores improved by roughly 20–25 units relative to random, and ProteinMPNN score increased by ~0.25 units, reflecting stronger sequence plausibility and structural compatibility.

Even at high similarity (80–90%), both methods performed significantly better than random for each metric (ESM-1v: *q* = 0.00036, *r* ≈ 1.0; ProteinMPNN: *q* = 0.00045, *r* ≈ 1.0).

Direct comparisons between the two masking strategies revealed minimal systematic differences: across most bins, variant-score distributions were statistically indistinguishable after multiple-testing correction (all *q >* 0.05). A modest advantage for APC emerged only in the 70–80% ESM-1v group (*q* = 0.0227, *r* = 0.56), potentially reflecting the benefit of attention-based residue coupling at moderate divergence.

Taken together, both sequence-based (ESM-1v) and structure-based (ProteinMPNN) metrics indicate that MSA-Transformer–guided pathway generation yields more biologically plausible intermediates relative to the random baseline.

### 2.3 Characterising Hybrid Sequences Through Sequence, Structure, and Latent Features

We next focus on characterising the intermediate sequences along the mutational pathways connecting the source (*S*) and target (*T*) proteins to identify potential hybrid variants. To quantify how intermediate sequences balance the features of both *S* and *T*, we define a hybrid score (*H*_sim_) that integrates sequence and structural similarity.

For a candidate sequence *C*, we compute its similarity to both *S* and *T* in sequence and structure space. Sequence similarity is measured using percent identity (Hamming similarity), while structural similarity is computed using structures predicted by ESMFold and compared with TM-align. The hybrid score is defined as

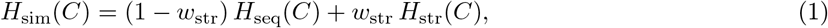

where *H*_seq_(*C*) = min {sim_seq_(*C, S*), sim_seq_(*C, T*) } and *H*_str_(*C*) = min sim_str_(*C, S*), {sim_str_(*C, T*) }. The weight *w*_str_ adapts the contribution of structure relative to sequence based on the similarity between the source and target proteins, increasing the importance of structural similarity at higher degrees of structural divergence. Further implementation details are provided in Supplementary Methods (Section S1.6).

Across pathways grouped by source–target sequence and structural similarity, a clear trend emerges (Figure 3). In the *low similarity* regime (10–60% sequence and 10–60% structure; Figure 3A), referred as Group A, intermediates are widely dispersed across the space of sequence and structural similarity, with most exhibiting low hybrid scores (*H*_sim_ *<* 0.6). At *moderate similarity* levels (60–80% for both sequence and structure; Figure 3B), corresponding to Group B, the distribution of *H*_sim_ values becomes more compact with a larger fraction of intermediates achieving hybrid scores above 0.8. When both sequence and structure are *highly similar* (≥80%; Figure 3C), representing Group C, most pathways yield hybrids with *H*_sim_ values in the 0.8–0.9 range. Because *H*_sim_ depends on the minimum similarity to both *S* and *T*, pairs with large sequence or structural divergence cannot achieve high hybrid scores, even if intermediates optimally bridge the two endpoints.

**Fig. 3.**
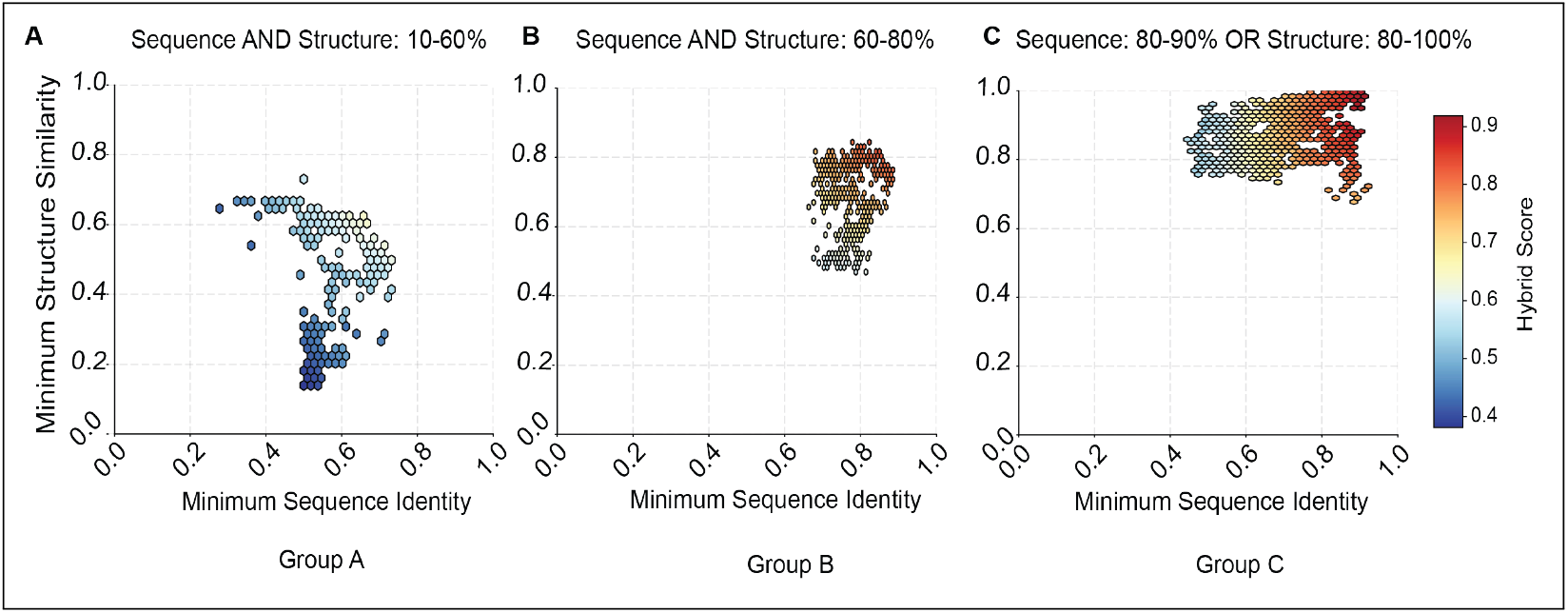
Hybrid scores characterise intermediate sequences. (A–C) Density plots (using hexagonal bins) showing the distribution of minimum sequence identity and minimum structural similarity for all intermediate sequences, coloured by hybrid score (*H*_sim_). Panels are grouped by source–target similarity ranges: (A) Group A: low similarity (10–60% sequence and 10–60% structure), (B) Group B: moderate similarity (60–80% sequence and structure), and (C) Group C: high similarity (*≥* 80% sequence or structure). Each hexagon represents the density of intermediates within that region of similarity space, with warmer colours corresponding to higher hybrid scores.

To further investigate the structural characteristics of the generated hybrids, representative intermediates from Groups A and B were selected for structural visualisation, chosen for their moderate to high *H*_sim_ values (Figure 3) and high in-silico variant scores (Result Section 2.2). Selected examples of hybrid structures are visualised and discussed in Figure 4.

**Fig. 4.**
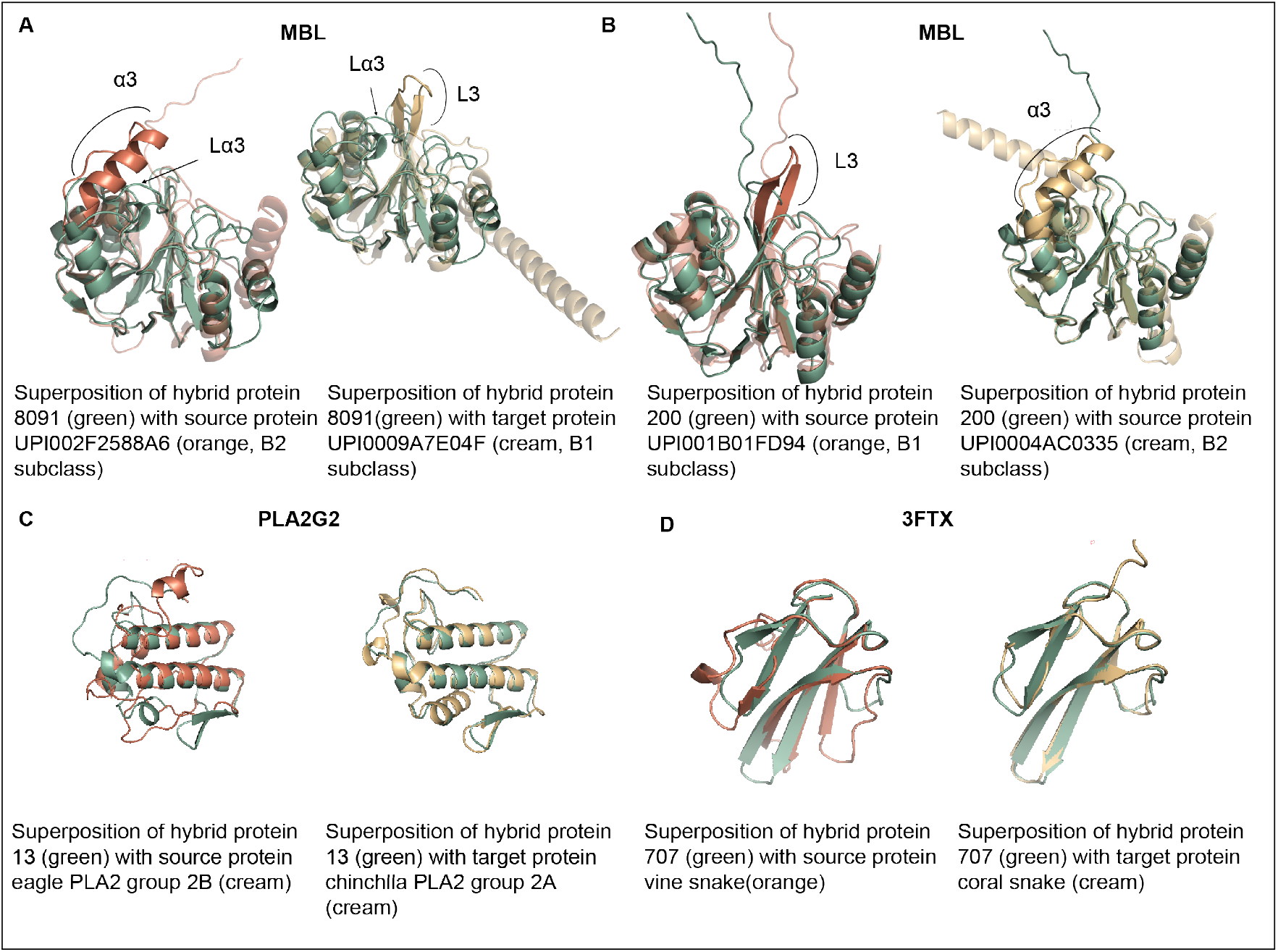
Example hybrid structures predicted using Chai [9] / ESMFold [22] and visualised in PyMOL [34]. (A) Hybrid 8091 derived from source protein UPI002F2588A6 (B2 subclass) and target protein UPI0009A7E04F (B1 subclass) from the MBLs. Hybrid 8091 contains a shortened L3 loop and lacks an extended *α*3 helix, resembling B2 and B1 MBLs respectively. The addition of a flexible loop, absent in both B1 and B2 MBLs, (L*α*3) may represent an alternative mode of substrate interaction. (B) Hybrid H200 derived from source protein UPI001B01FD94 (B1 subclass) and target protein UPI0004AC0335 (B2 subclass) from the MBLs. Hybrid 200 displays a novel combination of L3 and *α*3 segments similar to Hybrid 8091. (C) Hybrid 13 was derived from a source PLA2 of the bird Aquila chrysaetos (subfamily G2B; bAqchB) and a target PLA2 of the mammal Chinchilla lanigera (subfamily G2A; mChlaA4). The hybrid displays *β*-sheets characteristic of the mammalian G2A subfamily, while lacking the small helices that distinguish the source and target forms. Nevertheless, the canonical PLA2G2 scaffold is preserved in the hybrid, including the compact and well-defined catalytic center. (D) Hybrid 707 was derived from a source long-chain 3FTX of the vine snake (Ahaetulla prasina, TR|A0A193CHL5|Ahaetulla_prasina), representing a plesiotypic state of the three-finger toxin family, and a source short-chain 3FTX from the coral snake (Micrurus surinamensis, NCBI|MICSU_IACN01100666.1_C226_26|Micrurus_surinamensis). The hybrid lacks the small helix present on finger III of the vine snake toxin, but notably extends the *β*-sheets of finger I, effectively “bridging” the *β*-structures of the two parental toxins. Interestingly, similar configurations can be found among naturally occurring 3FTXs.

Among the MBLs, several hybrid candidates exhibited novel combinations of sub-class-specific structural elements, where one sub-class (e.g., B1) served as the source and the other (e.g., B2) as the target. In many such examples, the novel combination involved variation in the L3 loop and *α*3 helix, elements which have previously been implicated in substrate specificity differences between the two sub-classes [13, 30]. Figure 4A and 4B highlights two hybrid candidates which contain a short L3 and lack an extended *α*3, hallmarks of the B2 and B1 sub-classes respectively. Although this combination results in a lack of substrate-stabilising residues of either class, hybrid protein 200 (Figure 4B) retains a tyrosine in its L3 loop that could plausibly interact with *β*-lactam substrates in a manner similar to that observed for aromatic residues in the L3 loop of B1 MBLs. Intriguingly, hybrid protein 8091 (Figure 4A), which displays the same novel combination of *α*3 and L3 elements as the former, is also predicted to contain a long, disordered loop inserted directly downstream of a short *α*3. While such a loop is not observed in any B1 or B2 MBLs, similarly disordered loops are present at this position in representatives of other MBL-fold sub-families, including the convergently evolved B3 MBLs [42, 21]. In hybrid protein 8091, this loop may provide an alternative substrate binding capabilities to that of either the source or target. This may suggest that MSA-Transformer leverages general learnt representations to compensate for changes arising from the loss or gain of features along the pathway between source and target. The hybrid sequences are listed in the Supplementary Data, Section S4.1.

While the vast majority of hybrid candidates from MBLs dataset were predicted to adopt the core MBL-fold, some demonstrated unusual structural variation not observed in either the source or target protein sequences. These variations are likely to impact overall stability or favourable substrate interactions (see Supplementary Figure S4 for representative examples).

For Group C, hybrid candidates were validated using pretrained sparse autoencoder (SAE) InterProt [1] based latent features to test whether the intermediates combine attributes of both *S* and *T*. Because sequences in this group exhibit minimal structural divergence, SAE latent feature analysis provides a complementary view that captures feature-level integration beyond sequence or structure alone.

Each SAE latent feature encodes a distinct concept of the protein representation space, some align with known biological features while others remain as yet uncharacterised. For example, SAE latent feature 2375 captures bipartite nuclear localization signals (NLS), which are specific sequence motifs generally following the pattern R/K(X)_10–12_KRXK that enable proteins to enter the nucleus. Although the biological meaning of many SAE latent features remains unknown, the SAE framework provides a mean to monitor how the abstract features are gained or lost across intermediate sequences generated between a source (*S*) and a target (*T*).

To assess how latent feature composition differs in individual intermediate sequences, we grouped the sparse autoencoder features into common, source-only, and target-only categories (see Supplementary Methods, Section S1.7). For each category, we quantified the percentage change in activation value relative to the source sequence across all intermediates. Among Group C source-target pair, common features generally showed stable or modestly increasing activation, while source-only features tended to decrease and target-only features increased relative to the source sequence (See Supplementary Figure S5; representative examples of two hybrid sequences are shown in the Figure 5).

**Fig. 5.**
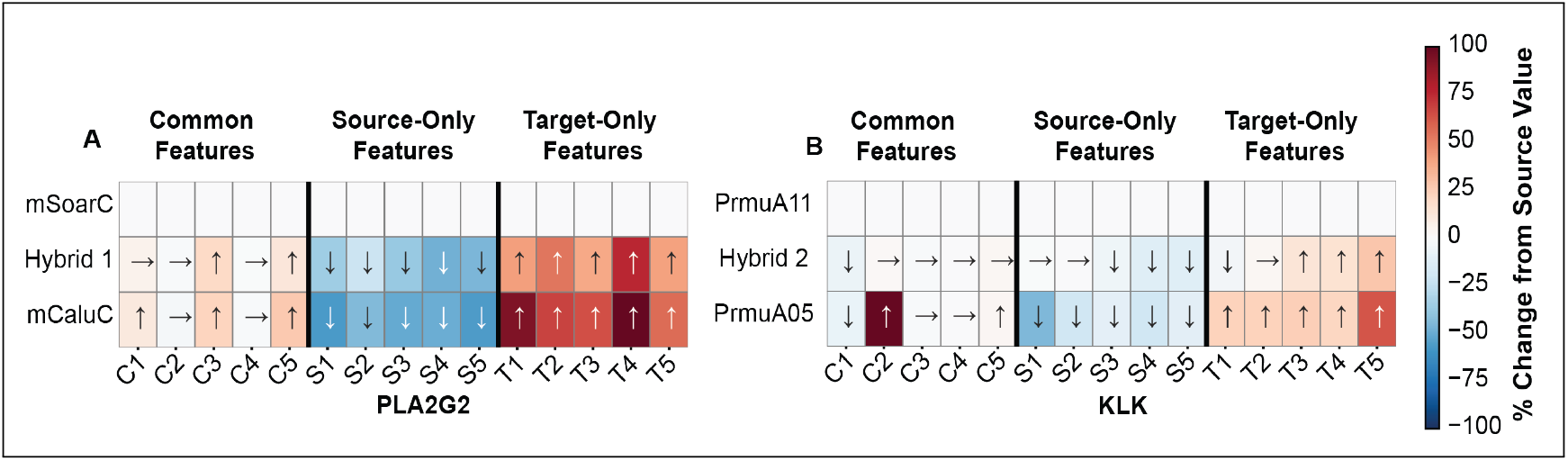
Hybrid sequences with high source–target similarity (Group C) display systematic shifts in SAE latent features relative to the source sequence. Panels (A) PLA2G2 and (B) KLK show heatmaps of percentage change in the sparse autoencoder (SAE) latent features relative to the source sequence. Features are grouped into common, source-only, and target-only categories (see Supplementary Methods, Section S1.7). Arrows indicate the direction of change, and colour intensity reflects the magnitude of deviation from the source value. In both examples, the hybrids show stable activation of common features, a reduction in source-only features, and an increase in target-only features—reflecting a gradual shift in latent feature composition.

## 3. Discussion

In this work, we introduced a protein sequence generator that steers the MSA-Transformer protein language model using the input MSA to generate intermediate variants *between* a source and a target protein within the same protein family. We generated these mutational pathways using two distinct masking strategies. Intermediate “hybrids” were evaluated by the extent to which they reflected or blended qualities of the source and target, based on similarities in sequence, structure and latent representations. Integrating these strategies with experimental validation could advance the design of hybrid proteins that combine desirable properties from multiple family members, bridging the gap between generative protein design and functional protein engineering.

The success of pathway generation depends on both the composition of the MSA and the sequence identity between the source and the target. We found MSAs containing between 15 to 90 sequences resembling the target sequence in embedding space (with a median intra-cluster cosine distance ranging from ~0.006 −0.05) were effective. Moderate sequence identity (60-70%) between source and target resulted in the highest convergence rate across both masking schemes (embedding-based and row-attention-guided) under our simulated annealing–based iterative framework. Across all similarity ranges, APC consistently achieved slightly higher convergence rates and required fewer iterations to reach convergence compared with IRS. Beyond these factors, the approach is also limited by the MSA-Transformer’s input length (1024 residues) and by GPU memory required by the model.

In this study, our ablations focused on the MSA conditioning context and the residue masking strategies (IRS and APC), while other optimisation parameters, such as the embedding distance metric, masking rate, and beam search configuration, were held fixed across experiments to isolate the effect of these design choices. A systematic exploration of these additional hyperparameters remains an interesting direction for future work.

Analysis of pathway geometry in the ESM2 model’s representation space revealed that successful pathways generated by the MSA-Transformer did not follow a straight line between the source and target embeddings. Instead, the pathways traced non-linear routes through the representation manifold, suggesting the model explores locally coherent regions rather than interpolating linearly between endpoints. This observation agrees with prior work showing that linear interpolation does not necessarily follow the curved geometry of learnt representation manifolds [40, 4]. Establishing biological baselines for pathway curvature, for example by analysing embedding-space transitions between naturally evolving orthologous proteins, would provide a useful reference for interpreting such deviations and represents an interesting direction for future work. Consistent with this interpretation, both sequence-based (ESM-1v) and structure-based (ProteinMPNN) variant scores indicated that the sequences generated along these pathways are significantly more plausible than those produced by random sampling, which followed a more linear pathway between source and target sequence.

While the present framework does not guarantee biologically viable sequences, it offers a generative mechanism that captures the directionality and continuity of mutational change within the model’s learned manifolds. Structural analysis of representative hybrids supports this notion: in the MBL family, hybrids combined subclass-specific motifs such as the shortened L3 loop of B2s and the lack of an extended *α*3 helix as found in B1s, and in some cases introduced novel flexible elements not present in either parent, suggesting plausible, non-canonical substrate-interaction modes. Similarly, hybrids in the PLA2G2 and 3FTX families preserved their canonical folds while recombining local features from source and target forms—for instance, the *β*-sheet arrangement of mammalian G2A PLA2s or the extended finger-I *β*-strands characteristic of short-chain 3FTSx. These examples illustrate that the model can produce structurally coherent yet compositionally mixed proteins, capturing non-trivial combinations of source and target features that may represent viable starting points for experimental characterisation or directed evolution.

To assess whether the generated sequences exhibit representation-level evidence of hybridisation, we analysed latent activations from a pre-trained sparse autoencoder (SAE) applied to sequence embeddings. This analysis probes how characteristics associated with the source and target sequences are represented along the mutational pathway. The biological concept captured by individual SAE latent features is not always known: while some correspond to recognisable properties such as secondary-structure motifs or specific functional domains, many capture abstract patterns whose biological interpretation remains unknown. Consequently, these activations should be viewed as representation-level diagnostics rather than direct indicators of specific biochemical attributes.

This study focuses on evaluating a specific hypothesis: whether an iterative framework combined with MSA-based protein language models capable of modelling insertions and deletions can generate intermediate hybrid proteins between related family members, and how such hybrids can be characterised. Because standardised benchmarks for generating hybrid proteins between predefined source and target sequences remain limited, we evaluate our framework using a simple random baseline together with widely used variant-scoring metrics. Developing more comprehensive benchmarks for comparing generative strategies represents an important direction for future work. Such efforts could include systematic comparisons with other generative models, for example by integrating single-sequence protein language models or training models to operate within controlled levels of sequence similarity.

Future work combining SAE-derived features with experimental validation could help determine whether these representation-level hybrid signals correspond to functional blending. Methodologically, training models to propose mutations within controlled ranges of sequence similarity, or integrating single-sequence language models to complement MSA-based approaches, may broaden the applicability of the framework. Additional structural context from models such as Pairformer [2] could further refine the characterisation of mutational landscapes. Developing reinforcement-learning strategies to select mutation sites, guided by the model’s predictions, also presents an opportunity to improve pathway efficiency and plausibility. Together, these avenues could enhance hybrid protein design and expand the exploration of the functional sequence space.

## Acknowledgments

We thank Dr Gabriel Foley for curating the KARI sequences. This work was supported by Australian Research Council Discovery Project 210101802 and National Health & Medical Research Council Ideas Project 2003871.

