## Supplementary for "Generating Hybrid Proteins with the MSA-Transformer"

for

#### S1 Supplementary Methods

##### S1.1 MSA-Transformer

The MSA-Transformer [9] is a protein language model trained in an unsupervised manner on 26 million multiple sequence alignments (MSAs), comprising on average ∼1,200 sequences each. It interleaves row and column attention to capture both inter-residue and inter-sequence dependencies, using a masked language-modeling objective that reconstructs masked amino acids from the surrounding MSA context. During training, a fraction (typically 15%) of amino acid positions in the MSA are masked uniformly at random, and the model is trained to reconstruct the true amino acids from the surrounding MSA context:

$$\mathcal{L}_{\mathrm{MLM}}(x;\theta) = -\sum_{(m,i)\in\mathrm{mask}} \log p(x_{mi} \mid \tilde{x};\theta) \tag{S1}$$

where $x_{mi}$ denotes the true amino acid at position $i$ in sequence $m$, and $\tilde{x}$ represents the masked MSA input. For each masked site, the model outputs a softmax-normalised probability distribution $p(x_{mi} \mid \tilde{x};\theta)$ over the 29-token vocabulary.

During inference, masked residues can be predicted either by sampling from the model's output distribution or by taking the maximum a posteriori (MAP) estimate:

$$\hat{x}_{mi} = \begin{cases} \text{sample from } \mathrm{Cat}(p(x_{mi} \mid \tilde{x};\theta)), & \text{(stochastic decoding)} \\ \arg\max_{a\in\mathcal{V}}\ p(a \mid \tilde{x};\theta), & \text{(deterministic decoding)} \end{cases} \tag{S2}$$

where $\mathcal{V}$ denotes the 29-token vocabulary including amino acids and other tokens.

Because the MSA Transformer operates under a masked language modelling formulation on aligned MSAs, predictions are made at existing alignment positions. Consequently, while the vocabulary includes gap tokens, the length of the alignment remains fixed and the model predicts tokens at each masked position without altering the overall aligned sequence length.

In addition to producing an output probability distribution over the vocabulary tokens, the MSA Transformer also produces contextualised representations for each position in the input MSA $M$. For a sequence of length $L$, the model outputs a representation tensor of shape $(L+1, d)$, where the first position corresponds to the CLS token and $d = 768$ is the embedding dimension. In this work, we discard the CLS token and obtain a sequence-level representation by mean pooling the position-wise representations across positions in the sequence. These pooled representations are subsequently used to compute sequence-level cosine distances between sequences and to guide the optimisation procedure described in Section S1.2.

##### S1.2 Prompting sequence changes from $S$ to $T$

To generate mutational pathways between a source sequence $S$ and a target sequence $T$, we leverage the MSA Transformer's probabilistic inference over masked amino acid sites (see Equation S2). The curated MSA $N$ (conditioning context), together with $S$ and $T$, forms the composite input MSA $M$ provided to the model.

To construct the conditioning context $N$, we first identify regions of high sequence density by clustering the sequences in the MSA (using their embeddings from the MSA Transformer) with a combination of HDBSCAN [6] and $k$-nearest neighbors (KNN). For each dense cluster, we compute the mean embedding and designate the sequence closest to this mean as the target $T$, while the remaining sequences in the cluster constitute the context set $N$. In cases where clusters are excessively large or heterogeneous, we refine them using available sequence annotations. For example, within the three-finger toxin (3FTx) protein family, annotated categories such as "plesiotypic," "long-chain," and "short-chain" are used to partition clusters into smaller, homogeneous subsets. Each resulting cluster contained 15-90 sequences resembling the target sequence in embedding space with a median intra-cluster cosine distance ranging from $\sim 0.006-0.05$, providing a compact and well-defined conditioning context for model steering.

At each iteration, a subset of positions $\mathcal{S} \subseteq \{1, \ldots, L\}$ in $S$ is masked. We employ two masking strategies—*Independent Residue Sampling* (IRS) and *Attention Position Coupling* (APC)—which differ in how they select residue sites for mutation (see Section S1.3). The modified MSA is then provided as input to the model, which predicts a conditional distribution over amino acids at the masked sites. New residues are selected deterministically as the most probable amino acid at each site, forming a candidate sequence $C$:

$$C = \arg\max_{a \in \mathcal{V}} p_\theta(a \mid S_{\setminus \mathcal{S}}, T, N), \tag{S3}$$

where $p_\theta$ denotes the conditional distribution inferred by the MSA Transformer, $S_{\setminus \mathcal{S}}$ represents the source sequence with masked positions $\mathcal{S}$, and $C$ is the proposed candidate sequence. Because predictions are made at existing alignment positions of the input MSA $M$, the aligned length of the candidate sequence $C$ remains fixed and identical to that of the sequences in $M$, including the source sequence $S$ and target sequence $T$.

Each candidate $C$ is evaluated using an objective function based on its cosine distance to the target sequence $T$ in the model's embedding space:

$$E(C) = 1 - \frac{\langle f(C), f(T) \rangle}{\|f(C)\| \, \|f(T)\|}, \tag{S4}$$

where $f(\cdot)$ denotes the sequence embedding obtained from the MSA Transformer, and $\langle \cdot, \cdot \rangle$ is the dot product. Lower $E(C)$ values indicate higher similarity to the target sequence.

Specifically, $f(\cdot)$ corresponds to the mean-pooled representation of the residue embeddings produced by the final layer of the MSA Transformer after excluding the CLS token. This pooling operation yields a fixed-length vector in $\mathbb{R}^{768}$, allowing cosine distance comparisons between the source, candidate, and target sequences.

Cosine distance is used to compare sequence embeddings, as cosine similarity is a standard metric for measuring representational similarity in embedding spaces and is commonly used in both natural language [7] and protein language models [8] In preliminary experiments, cosine distance resulted in more consistent convergence of mutational trajectories compared to Euclidean distance. Exploring alternative distance metrics remains an interesting direction for future work.

Candidate sequences are then accepted probabilistically according to a simulated annealing criterion:

$$P_{\text{accept}} = \begin{cases} 1, & \text{if } \Delta E \leq 0, \\ \exp(-\Delta E/\tau), & \text{if } \Delta E > 0, \end{cases} \tag{S5}$$

where $\Delta E = E(C) - E_{\text{current}}$ is the change in cosine distance, and $\tau$ is the temperature parameter. We use $\tau$ as 1 with decay of 0.99 for all our experiements. Early iterations use a high $\tau$ to encourage exploration of sequence space, while later iterations progressively reduce $\tau$ to focus on higher-quality candidates.

Each iteration comprises $t$ independent "tosses." For every toss, a candidate sequence is generated according to the selected masking strategy. After all $t$ tosses, a total of $c \cdot t$ candidates are obtained, from which the best sequence is chosen based on the annealing criterion and objective function. This multi-proposal scheme enhances exploration of the local sequence neighborhood while maintaining selection pressure toward optimal transitions.

The process continues until either a maximum number of iterations is reached or convergence is achieved. Convergence is defined as the point at which the cosine distance to the target sequence falls below a threshold $d_{\text{conv}} = \gamma \cdot d(S, T)$, where $\gamma$ is the convergence factor. For all experiments, we set $\gamma = 0.25$, corresponding to traversing 75% of the cosine distance between $S$ and $T$.

##### S1.3 Masking positions in $S$

We employ two masking strategies: *Independent Residue Sampling* (IRS) and *Attention Position Coupling* (APC). In both cases, a residue-wise masking probability profile is first constructed based on the cosine distance between the residue embeddings of $S$ and $T$. Positions with low entropy, computed from the model's output logits, are excluded from masking. We interpret low-entropy positions as structurally or functionally constrained sites that should be altered only after certain dependent residues have changed. To enforce this constraint, positions falling below the 30th entropy percentile are filtered out; this percentile can be treated as a tunable hyperparameter.

In the IRS strategy, sites are then chosen independently from this masking probability profile. In contrast, the APC strategy incorporates inter-residue dependencies captured by the MSA Transformer's attention maps. An average row-attention map is computed across all layers and heads to obtain a single residue–residue attention matrix describing coupling between alignment positions, followed by the application of average product correction to remove background correlation bias and emphasise meaningful couplings between positions. A seed position is first selected from the residue-wise masking probability profile, after which the corrected row-attention map is used to identify the $n_{\text{att}}$ positions with the highest mean attention to the sampled site. We set $n_{\text{att}} = 10$ for all experiments. These positions are then added to the mask set, and the procedure is repeated until the desired number of masked residues is reached.

##### S1.4 Enhancing generation of changes by beam search

To improve the diversity and quality of generated mutational pathways, we extend the iterative masking procedure (Section S1.2) with a beam search framework [11]. Rather than following a single path of sequence updates, multiple candidate sequences are maintained and evaluated in parallel at each iteration, forming a beam of width $W$ representing competing pathways. This approach enables the exploration of multiple routes between the source $S$ and target $T$.

Within the beam search, we employ stochastic decoding, in which amino acids at masked positions are sampled from the MSA Transformer's conditional probability distribution rather than selected deterministically. For each masked configuration, the top-$k$ candidates are drawn from this conditional distribution, allowing exploration of multiple plausible residue substitutions at each iteration. With beam width $W$ and $t$ tosses per iteration, this procedure yields a total of $W \cdot k \cdot t$ candidate sequences per iteration.

All candidates are evaluated using the cosine distance objective combined with the simulated annealing acceptance criterion (same as Section S1.2). Among the accepted candidates, sequences are ranked according to their log-likelihood under the MSA Transformer, and the top $W$ sequences are retained to form the next beam. A candidate exits the beam once it meets the convergence condition $d(C, T) \leq d_{\text{conv}}$. The process continues until either the maximum number of iterations is reached or no candidates remain in the beam.

Beam search simulations were run for 100 iterations with a masking rate of 5% of sequence length. For pairs that failed to converge after these runs, the masking rate was increased to 10% while maintaining the same iteration schedule to assess the effect of broader mutation coverage on convergence. The 5-10% masking rate was chosen to introduce incremental changes at each step. This rate is lower than or similar to the masking proportion used during the original MSA-Transformer training objective, ensuring that the model operates within the distributional regime it was trained on. Across all experiments, we use $t$ as 3, $W$ as 3 and $k$ as 5.

##### S1.5 Deviation Score

The deviation score measures the geometric deviation of the pathway from the direct (linear) interpolation connecting $S$ and $T$. Let $\mathbf{v} = \mathbf{E}_t - \mathbf{E}_s$ represent the directional vector from source to the target, where $\mathbf{E}_t$ and $\mathbf{E}_s$ are their representative embeddings. For each intermediate candidate sequence $Ci$ with embedding $\mathbf{E}_i$, we project $\mathbf{E}_i$ onto the line segment connecting $S$ and $T$:

$$\alpha = \frac{(\mathbf{E}_i - \mathbf{E}_s) \cdot \mathbf{v}}{\mathbf{v} \cdot \mathbf{v}}, \quad \mathbf{E}_\alpha = \mathbf{E}_s + \alpha \cdot \mathbf{v}$$

where $E_\alpha$ is the projected $E_i$ on $v$.

The *Deviation Score* is defined as the Euclidean distance between the actual embedding and its projection onto this line:

$$D_i = \|\mathbf{E}_i - \mathbf{E}_\alpha\|.$$

A smaller $D_i$ indicates that the pathway closely follows the linear path between $A$ and $B$, while a larger value signifies divergence from this direct route through embedding space.

##### S1.6 Hybrid Score

The Hybrid Scoring scheme quantifies the likeness of each sequence in the pathway to both the source ($S$) and target ($T$) sequences. We define hybridness using two complementary approaches: one based on **sequence and structural similarity**, and another based on **latent feature representations** derived from a pretrained sparse autoencoder [1].

In the first approach, we assess sequence and structural similarity of all intermediate sequences relative to the source and target. Sequence similarity is quantified using Hamming distance (percent identity), while structural similarity is evaluated using ESMFold [5] and TM-align [13]. These complementary measures capture how sequences progressively replace or mix source and target features along the pathway.

For each source–target pair, we compute the sequence similarity $\mathrm{sim}_{\mathrm{seq}}(S,T)$ and structural similarity $\mathrm{sim}_{\mathrm{str}}(S,T)$, which represent the overall similarity between the source ($S$) and target ($T$) sequences. These values capture how closely related the two proteins are in sequence and structure, and are used to define a fixed structural weight that balances the relative contribution of structural and sequence information:

$$w_{\mathrm{str}} = \frac{1 - \mathrm{sim}_{\mathrm{str}}(S,T)}{(1 - \mathrm{sim}_{\mathrm{str}}(S,T)) + (1 - \mathrm{sim}_{\mathrm{seq}}(S,T))}. \tag{S6}$$

For a candidate sequence $C$, the overall hybrid score $H_{\mathrm{sim}}(C)$ is computed as a weighted combination of sequence- and structure-based similarity:

$$H_{\mathrm{sim}}(C) = (1 - w_{\mathrm{str}})\, H_{\mathrm{seq}}(C) + w_{\mathrm{str}}\, H_{\mathrm{str}}(C), \tag{S7}$$

where the per-channel similarity is defined as

$$H_{\mathrm{seq}}(C) = \min\{\mathrm{sim}_{\mathrm{seq}}(C,S),\, \mathrm{sim}_{\mathrm{seq}}(C,T)\}, \qquad H_{\mathrm{str}}(C) = \min\{\mathrm{sim}_{\mathrm{str}}(C,S),\, \mathrm{sim}_{\mathrm{str}}(C,T)\}. \tag{S8}$$

This formulation adaptively emphasises structural similarity when the source and target structures differ substantially, while prioritizing sequence similarity when the structures are nearly identical.

##### S1.7 Sparse Autoencoder–Derived Latent Features–Based Evaluation of Hybrid Sequences

To capture and evaluate hybrid properties encompassing, but not limited to, sequence and structural similarity, we incorporated latent features derived from the pretrained sparse autoencoder (SAE) InterProt [1]. Each protein sequence was encoded into a 4096-dimensional SAE latent vector, representing the mean activation across all sequence positions.

To focus on the most informative SAE latent features, we first excluded features with mean activation value below 0.1 across the dataset. For each source–target pair, we then identified the top 500 most active features separately for the source ($S$) and target ($T$) sequences based on their mean activation values. The union of these top features was partitioned into three categories:

– Source-only features, present among the top 500 in $S$ but not in $T$;
– Target-only features, present among the top 500 in $T$ but not in $S$; and
– Common features, shared between $S$ and $T$, further ranked by their variance across intermediates.

Each intermediate sequence was then evaluated based on the relative activation of these SAE latent feature sets, allowing us to trace how source- and target-associated latent features emerge, diminish, or co-occur along the mutational pathway.

##### S1.8 Datasets

We selected five protein families for the experiments performed in this study: Ketol-acid reductoisomerase (KARI) Class I, Phospholipasea A2 group 2 (PLA2G2), B1/B2 lactamases (MBLs), Three-finger toxins (3FTx), and Tissue Kallikreins (KLK). Families were selected to span diverse folds, functions, and evolutionary histories. For the Pla2g2, 3FTx and KLK families, we combined toxic and non-toxic homologs, sampling both derived and ancestral-like states to capture broad evolutionary variation during emergence and diversification of novel functions.

For 3FTx, we used the previously published dataset [4], which integrates toxic 3FTx genes from caenophidian snakes with non-toxic uPAR/LY6 family homologs across vertebrates, capturing both the toxin radiation and ancestral non-toxic states. The KLK dataset is from Barua et al. [2] that jointly analyses mammalian and reptilian (including snake) kallikreins, sampling canonical tissue KLKs alongside independently co-opted venom KLK-like toxins, thereby spanning non-toxic and derived toxic functions across vertebrates.

For Pla2g2, we used the previously compiled PLA2G2 dataset of Koludarov et al. [3], which delineates the g2 locus across 93 vertebrate genomes and documents multiple co-options into venom, alongside innate-immunity and other non-toxic roles, thus covering both ancestral and derived states across tetrapods. For the lactamases (MBLs) family, sequences for B1 and B2 MBLs were curated using clade-specific profile Hidden Markov Models (pHMMs). Hits from the Uniparc database were clustered at 90% sequence identity using the cluster program of the MMseqs2 package, prior to alignment with MAFFT (v7.487 with the localpair option). B1 and B2 MBLs were identified on the basis of known subgroup-specific metal-binding residues. In order to constrain the alignment to a length amenable to MSA Transformer, sequences with a high proportion of content in low-occupancy columns were removed. This yielded 1622 B1 MBLs and 33 B2 MBLs. Subsequently, in order to decrease sequence volume while maintaining diversity, B1 MBL sequences were clustered at 80% sequence identity and further subset by sampling 617 cluster representatives.

For KARI protein family, sequences used have an EC number of either 1.1.1.86, 1.1.1.382, or 1.1.1.383, representing KARIs that are NADP+-, NAD+- or non- specific. This initial sequence dataset was condensed by clustering using CD-HIT at 70% identity followed by the removal of sequences with non-canonical amino acids, sequences that are either less than 250 and greater than 400 amino acids in length, and sequences annotated by UniProt as containing disordered regions, fragments, and non-terminal residues. Furthermore, family-specific properties were also considered in selecting sequences. These properties include the presence of the GxGxxG motif [12], as well as the UniPathway tags UPA00047 (L- isoleucine biosynthesis) and UPA00049 (L-valine biosynthesis). The selected sequences also only contain the CATH superfamily tags 6.10.240.10 and 3.40.50.720 (comprising the NAD(P)-binding Rossmann-like domain), and only the PROSITE tags PS51850 (KARI N- terminal domain) and PS51851 (KARI C-terminal domain). By filtering sequences using information from a range of structural and evolutionary databases, the diversity of sequences was maximised without including KARIs that might be non-functional or misannotated in UniProt. The final dataset consisted of 716 sequences, which were aligned to identify homologous positions using MAFFT-DASH [10].

#### S2 Supplementary Figures

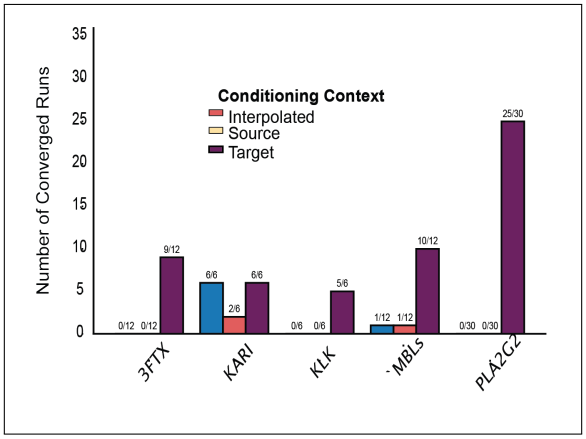


**Fig. S1.** Convergence of sequences toward target sequences under different MSA conditioning contexts for masking strategy IRS across multiple protein families.

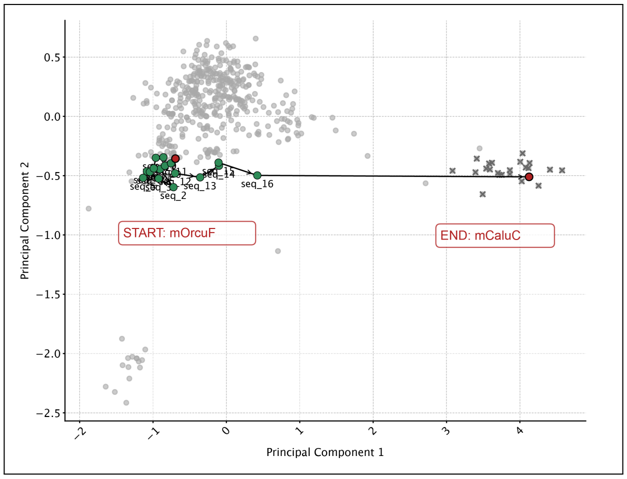


**Fig. S2.** Mutational pathway from $S$ ($mOrcuF$) to $T$ ($mCaluc$) within the PLA2G2 family, visualised in PCA space with IRS masking strategy.

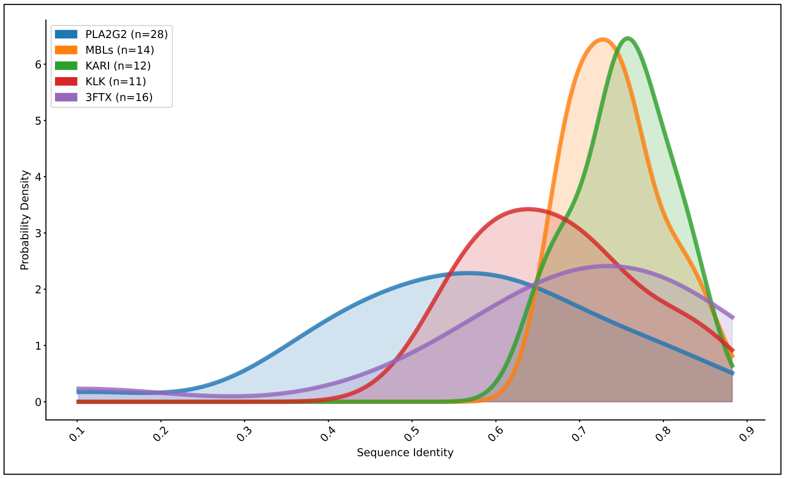


**Fig. S3.** Kernel density distribution of source–target sequence pairs used for mutational pathway generation, coloured by protein family. The plot shows that the selected sequence pairs span a broad range of sequence identities across five representative protein families (PLA2G2, MBLs, KARI, KLK, and 3FTx).

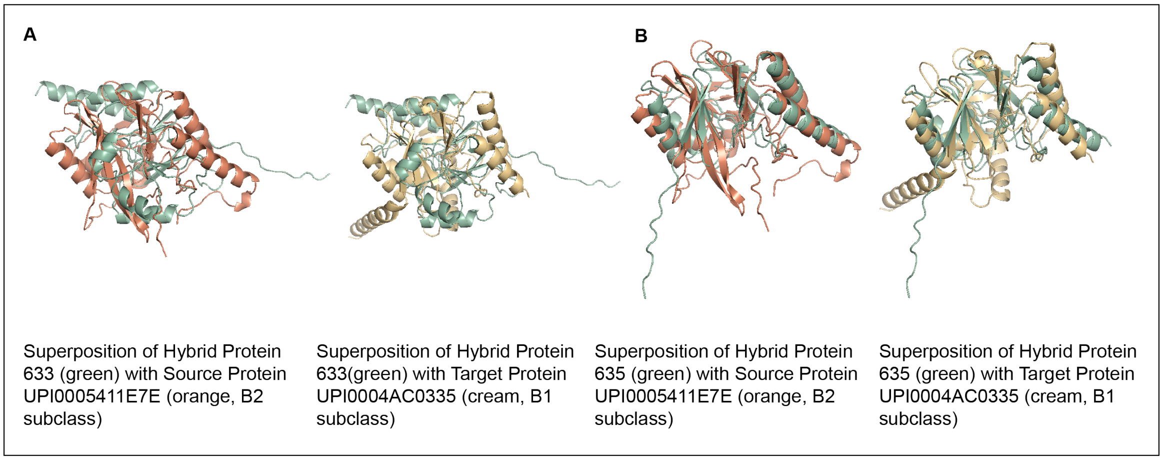


**Fig. S4.** Example Hybrid MBLs with unusual structural variation not observed in either the source or target protein sequences. These variations are likely to impact overall stability or favourable substrate interactions.

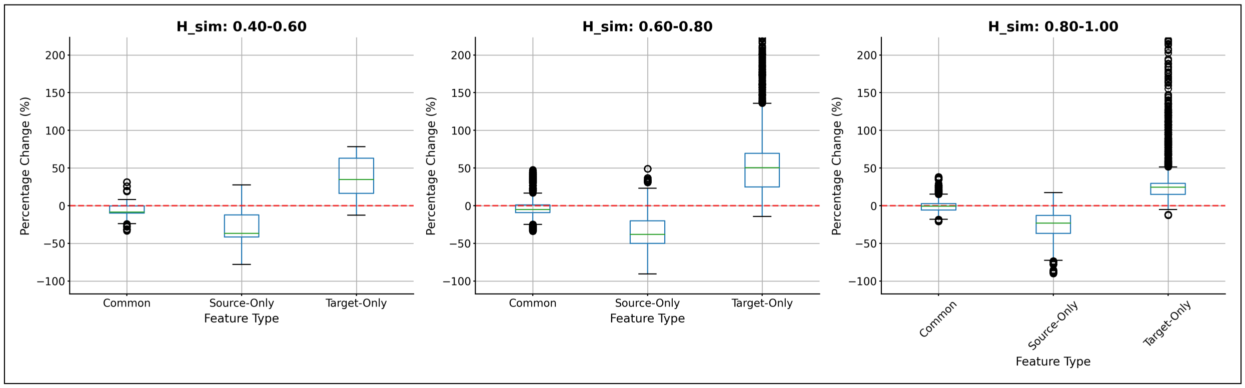


**Fig. S5.** Distribution of percentage change in SAE latent feature values across all intermediate sequences for the Group C source–target pairs. Boxplots show percentage change in activation for common, source-only, and target-only latent features relative to the source sequence, grouped by hybrid similarity bins ($H_{\mathrm{sim}}$). The red dashed line indicates no change relative to the source. At high $H_{\mathrm{sim}}$ values, the relative changes in source- and target-only features become more balanced.

#### S3 Supplementary Tables

**Table S1.** Summary of total convergence counts for the IRS and APC masking strategy under varying masking percentages for mutational pathway generation.

| Identity | 5% Masking | | 10% Masking | | Total Converged | |
|---|---|---|---|---|---|---|
| | **IRS** | **APC** | **IRS** | **APC** | **IRS** | **APC** |
| 10–50% | 6 | 7 | 1 | 1 | 7/13 | 8/13 |
| 50–60% | 5 | 6 | 3 | 4 | 8/10 | 10/10 |
| 60–70% | 18 | 18 | 1 | 2 | 19/20 | 20/20 |
| 70–80% | 19 | 19 | 1 | 2 | 20/23 | 21/23 |
| 80–90% | 8 | 8 | 5 | 5 | 13/15 | 13/15 |

**Table S2.** Average number of iterations required for convergence by the IRS and APC masking strategy across different masking percentages.

| Identity | 5% Masking | | 10% Masking | | Average Iterations | |
|---|---|---|---|---|---|---|
| | **IRS** | **APC** | **IRS** | **APC** | **IRS** | **APC** |
| 10–50% | 43 | 20 | 30 | 70 | 41 | 26 |
| 50–60% | 15 | 15 | 18 | 13 | 16 | 14 |
| 60–70% | 18 | 13 | 7 | 54 | 17 | 17 |
| 70–80% | 16 | 11 | 7 | 8 | 15 | 10 |
| 80–90% | 8 | 6 | 15 | 10 | 11 | 7 |

**Table S3.** Summary of paired Wilcoxon signed-rank test results for deviation scores across sequence identity bins. Values represent mean (standard deviation) for each method. P-values were adjusted for multiple comparisons using the Benjamini–Hochberg false discovery rate (FDR) procedure.

| Identity | Comparison | n_pairs | Random | APC / IRS | p-value | FDR q-value | Effect size ($r$) |
|---|---|---|---|---|---|---|---|
| 10–50% | APC vs Random | 8 | 0.57 (0.21) | 1.16 (0.66) | $0.0078^{**}$ | 0.0156 | 0.94 |
| | IRS vs Random | 7 | 0.56 (0.23) | 1.37 (0.76) | $0.0156^{*}$ | 0.0260 | 0.91 |
| 50–60% | APC vs Random | 10 | 0.62 (0.26) | 1.01 (0.56) | $0.0273^{*}$ | 0.0390 | 0.70 |
| | IRS vs Random | 8 | 0.50 (0.10) | 1.08 (0.51) | $0.0156^{*}$ | 0.0260 | 0.85 |
| 60–70% | APC vs Random | 20 | 0.51 (0.17) | 0.83 (0.54) | $0.0010^{**}$ | 0.0017 | 0.73 |
| | IRS vs Random | 19 | 0.50 (0.16) | 0.93 (0.47) | $0.000019^{***}$ | 0.000063 | 0.98 |
| 70–80% | APC vs Random | 21 | 0.37 (0.11) | 0.72 (0.32) | $0.000031^{***}$ | 0.000078 | 0.91 |
| | IRS vs Random | 20 | 0.38 (0.10) | 0.87 (0.42) | $0.000013^{***}$ | 0.000065 | 0.97 |
| 80–90% | APC vs Random | 13 | 0.23 (0.07) | 0.41 (0.16) | $0.00024^{***}$ | 0.00048 | 1.00 |
| | IRS vs Random | 13 | 0.23 (0.07) | 0.43 (0.17) | $0.00049^{***}$ | 0.00082 | 0.97 |

**Table S4.** Summary of paired Wilcoxon signed-rank test results for ESM-1v variant scores across sequence identity bins. Values represent mean (standard deviation) for each method. P-values were adjusted for multiple comparisons using the Benjamini–Hochberg false discovery rate (FDR) procedure.

| Identity | Comparison | n_pairs | Random | APC / IRS | p-value | FDR q-value | Effect size ($r$) |
|---|---|---|---|---|---|---|---|
| 10–50% | APC vs Random | 8 | -38.90 (6.05) | -28.80 (7.72) | 0.0156* | 0.0260 | 0.85 |
| | IRS vs Random | 7 | -37.36 (4.79) | -26.11 (7.60) | 0.0156* | 0.0260 | 0.91 |
| | APC vs IRS | 7 | – | – | 0.38 (ns) | 0.475 | 0.34 |
| 50–60% | APC vs Random | 10 | -47.05 (17.77) | -34.61 (17.08) | 0.0273* | 0.041 | 0.70 |
| | IRS vs Random | 8 | -46.34 (18.79) | -34.56 (16.06) | 0.055 (ns) | 0.073 | 0.68 |
| | APC vs IRS | 8 | – | – | 0.0156* | 0.0260 | 0.85 |
| 60–70% | APC vs Random | 20 | -57.49 (25.20) | -36.77 (16.43) | 0.000002*** | 0.0000075 | 1.00 |
| | IRS vs Random | 19 | -59.06 (24.89) | -33.27 (14.67) | 0.000019*** | 0.000041 | 0.98 |
| | APC vs IRS | 19 | – | – | 0.080 (ns) | 0.10 | 0.40 |
| 70–80% | APC vs Random | 21 | -59.52 (20.64) | -34.37 (16.11) | 0.000002*** | 0.0000075 | 1.00 |
| | IRS vs Random | 20 | -60.97 (20.08) | -32.22 (15.01) | 0.000002*** | 0.0000075 | 1.00 |
| | APC vs IRS | 20 | – | – | 0.0121* | 0.0227 | 0.56 |
| 80–90% | APC vs Random | 13 | -54.11 (18.73) | -38.02 (17.03) | 0.00024*** | 0.00036 | 1.00 |
| | IRS vs Random | 13 | -54.11 (18.73) | -38.19 (17.19) | 0.00024*** | 0.00036 | 1.00 |
| | APC vs IRS | 13 | – | – | 0.84 (ns) | 0.84 | 0.06 |

**Table S5.** Summary of paired Wilcoxon signed-rank test results for ProteinMPNN variant scores across sequence identity bins. Values represent mean (standard deviation) for each method. P-values were adjusted for multiple comparisons using the Benjamini–Hochberg false discovery rate (FDR) procedure.

| Identity | Comparison | n_pairs | Random | APC / IRS | p-value | FDR q-value | Effect size ($r$) |
|---|---|---|---|---|---|---|---|
| 10–50% | APC vs Random | 8 | -1.77 (0.11) | -1.64 (0.12) | 0.0078** | 0.0167 | 0.94 |
| | IRS vs Random | 7 | -1.76 (0.12) | -1.52 (0.12) | 0.0156* | 0.0260 | 0.91 |
| | APC vs IRS | 7 | – | – | 0.22 (ns) | 0.33 | 0.46 |
| 50–60% | APC vs Random | 10 | -1.76 (0.21) | -1.53 (0.15) | 0.0195* | 0.0293 | 0.74 |
| | IRS vs Random | 8 | -1.71 (0.16) | -1.58 (0.11) | 0.078 (ns) | 0.104 | 0.62 |
| | APC vs IRS | 8 | – | – | 0.74 (ns) | 0.89 | 0.12 |
| 60–70% | APC vs Random | 20 | -1.78 (0.19) | -1.54 (0.19) | 0.000004*** | 0.000015 | 1.00 |
| | IRS vs Random | 19 | -1.77 (0.19) | -1.55 (0.22) | 0.000027*** | 0.000068 | 0.96 |
| | APC vs IRS | 19 | – | – | 0.49 (ns) | 0.63 | 0.16 |
| 70–80% | APC vs Random | 21 | -1.75 (0.14) | -1.53 (0.13) | 0.000001*** | 0.000015 | 1.00 |
| | IRS vs Random | 20 | -1.76 (0.14) | -1.51 (0.14) | 0.000002*** | 0.000015 | 1.00 |
| | APC vs IRS | 20 | – | – | 0.28 (ns) | 0.39 | 0.24 |
| 80–90% | APC vs Random | 13 | -1.72 (0.15) | -1.57 (0.14) | 0.00024*** | 0.00045 | 1.00 |
| | IRS vs Random | 13 | -1.72 (0.15) | -1.56 (0.14) | 0.00024*** | 0.00045 | 1.00 |
| | APC vs IRS | 13 | – | – | 0.95 (ns) | 0.95 | 0.02 |

### S4 Supplementary Data

#### S4.1 MBL Protein Sequences

```
>UPI0005411E7E
MVGKCALFSFFLASLSLSAADSQLTITPLSEGVYEHISYQQVGKWGNVAARGLVVVDD
KDAYIIDTPWSNDDTLKLVEWAKDQGFMLKAAVVTHFHQDASGGLDVLNKRNIPTYAYQ
ETNRLLKQHKGLSAKHTIEETPFSLLQDKIDVFYPGGGHTADNVVVWLEQHQMLFAGCF
VKGLNSKHLGNLEDAVVAQWPTSIENTLNTYPTIKQVFPGHGRSGDQSLLLHTAKLAEN
YLAEQKSNPSSGSILKAH
>UPI0004AC0335
MKLMYVLMVSLTLIALNSVAHEGAITLTHFKGPLYIVEDKAYVQENSMVYIGADDITII
GATWTPETAEALEKEIRKVSALPIKEVINTNYHTDRAGGNAHWKKLGASIVSTQMTYDL
EKSQWKSIVDFTRQGFDKYPRLKESLPDKVYSGDFELQSGRVRALYLGPAHTEDGIFVY
FPSERVLYGNCILKEKLGNLSFANRTEYPKTLKKLKELIDQGELQVESIIAGHDTPIHG
VELIDHYLALLEDAKK
>UPI002F2588A6
MVRLVCGSMLLTALAACGAAPEITLSHLRGPLYVAEDSYYTKENAMVYVGASFVTVIGA
TWTPETARLLAEEIRKVTPKPVKEVVNTNYHPDRAGGNAYFKSIGAKVVATQMTYDLLE
RNWASVVDWTRAAFPDYPRLPLVLPDVVHPGDFELQDGRIRTFYLGPSHTPDGIFVYFP
EEKVLYGGCILKERLGNLDFANLVEYPKTLARLKRLDLPITTIVAGHWSPLHGPELIDE
YLKLLDQHSHEQR
>UPI0009A7E04F
MKRLLILSVVLAITVATSAGITITNHLRLRYYTADSILTVYKSDNLIIRKLSRNVYEHT
SFLNTNTFGRVSCNGMIVSNKQEAVVFDTPSTNSDSEELLRFLINDLKLQVKAIVATHF
HADCLGGLDSFHAMGVKSYANVKTIALAKSVNATIPQKSFKDHFSFKVGNKSVHVEFLG
EGHTRDNVVAYFPDDKILFGGCLVKELNASKGNLADANVNAWPETIRKVYDKYPNTQIV
IPGHGEIGGLELLDYTRKLFK
>hybrid_8091
MLLGSMLLAAGELTLSHLRGPLYVADGYGENAMVYVDASGVVVIDAPWTPEQARLLAEE
LEKRGVPVKEVVVTHYHDDCLGGLEYFKSLGAKVYATQMTYDLLERGRPEWAFPDPPRR
RLVPDVVFPGDLELGIRTYYLGPGHTPDNIVVYFPEEKVLYGGCILKERGNLDDANLVE
YPKTLARLRRLGITTIVAGHGSPELIDETLKLLDQHHHE
>hybrid_200
LLFLCLCCLTTASLAAEAEITLTHLKGPLYVVEDSYYVKENSMVYIGEKYITVIGATWT
PETAKLLAKEIEQVSHQPIKEVINTNYHPDRAGGNAYWKSIGAKIVSTQMTYDLLKEPD
KVYPGDFELQDGKVKAFYLGPSHTPDGIFVYFPEEKVLYGNCILKEQLGNLDFANLEEY
PKTLQKLKSKNLDIKTIIAGHDSPIHGPELIDHYLQLLAK
```
